## supplemental material for "Molecular motion and tridimensional nanoscale localization of kindlin control integrin activation in focal adhesions"

* Co-last authors

**This part includes**

Supplementary figures 1 to 8

Supplementary tables 1 to 7

Supplementary movie legends 1 to 3

**Other Supplementary Information for this manuscript includes the following:**

Supplementary Movies 1 to 3

#### SUPPLEMENTARY FIGURES

### Figure S1

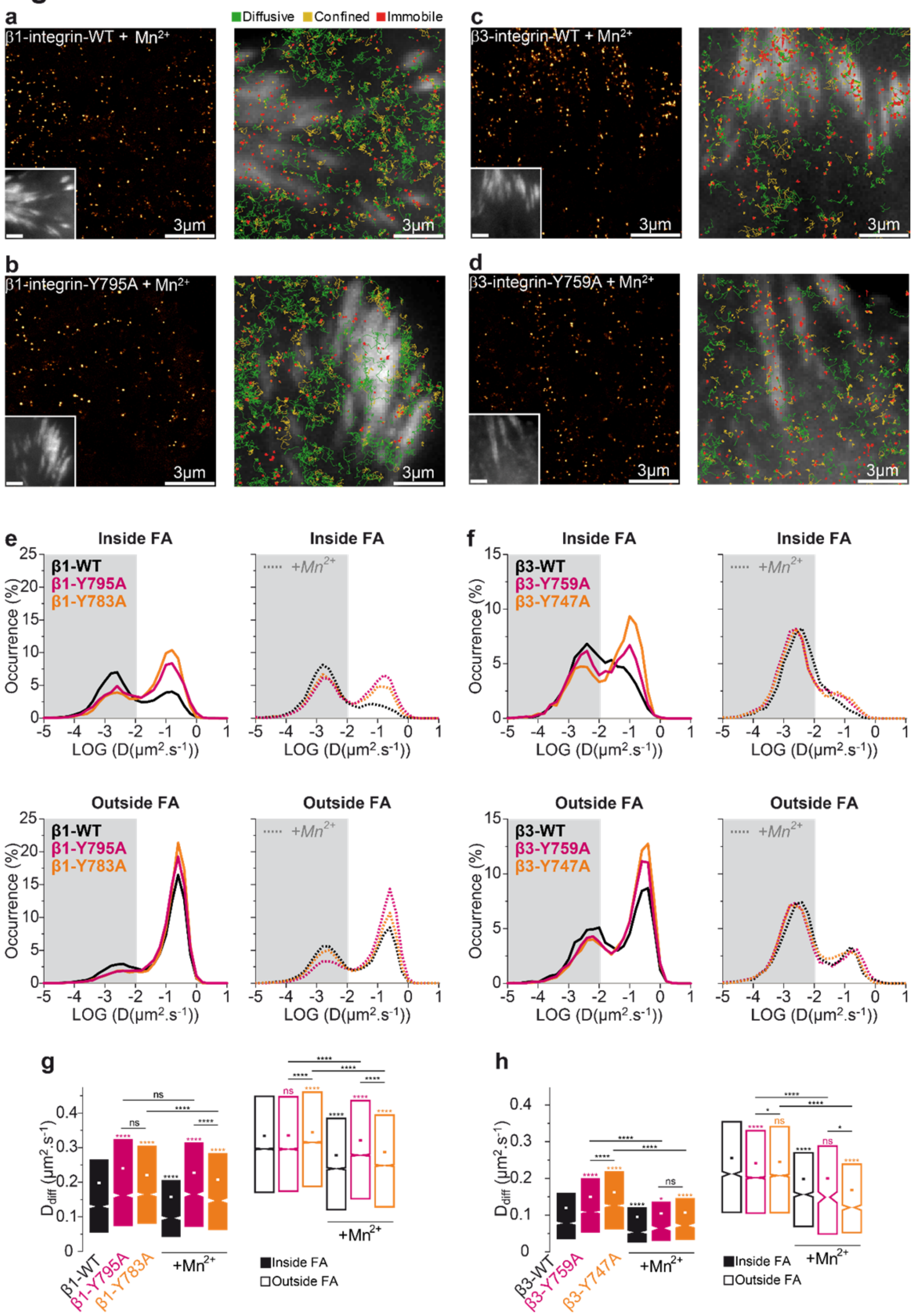

**Supplementary figure 1. Kindlin is required for $Mn^{2+}$-induced β1-integrin immobilization.** (**a**-**d**) Left: Super-resolution intensity images of β1-WT-mEos2 (a), β1-Y95A-mEos2 (b), β3-WT-mEos2 (c), β3-Y759A-mEos2 (d) in $Mn^{2+}$-stimulated MEFs obtained from a sptPALM sequence (50 Hz, >80 s). Inset: low resolution image of GFP-paxillin, which was co-expressed for FAs labelling (scale bar: 3 µm). Right: color-coded trajectories overlaid on FAs labelled by GFP-paxillin (greyscale) show the diffusion modes: free diffusion (green), confined diffusion (yellow) and immobilization (red). (**e**) Distributions of the diffusion coefficient D, computed from the trajectories obtained inside (top) and outside (bottom) FAs with β1-WT-mEos2 (black), β1-Y795A-mEos2 (pink), β1-Y783A-mEos2 (orange) in MEFs (left, full lines) and in $Mn^{2+}$-stimulated MEFs (right, dashed lines), are shown in a logarithmic scale. The grey area including D values inferior to 0.011 $µm^2.s^{-1}$ corresponds to immobilized proteins. Values represent the average of the distributions obtained from different cells. (**f**) Same as **e**, but with β3-WT-mEos2 (black), β3-Y759A-mEos2 (pink), β3-Y747A-mEos2 (orange). (**g**) Box plots displaying the median (notch) and mean (square) ± percentile (25–75%) of diffusion coefficients corresponding to the free diffusion trajectories of β1-WT-mEos2 (black), β1-Y795A-mEos2 (pink), β1-Y783A-mEos2 (orange) inside (left) and outside (right) FAs. (**h**) Same as **g**, but with β3-WT-mEos2 (black), β3-Y759A-mEos2 (pink), β3-Y747A-mEos2 (orange). Results for β1-WT-mEos2 without (16 cells) and with $Mn^{2+}$ (15 cells); β1-Y795A-mEos2 (22 cells) and with $Mn^{2+}$ (22 cells); β1-Y783A-mEos2 without (16 cells) and with $Mn^{2+}$ (20 cells); β3-mEos2 without (16 cells) and with $Mn^{2+}$ (13 cells); β3-Y759A-mEos2 without (20 cells) and with $Mn^{2+}$ (11 cells, 2 ind. exp.) and β3-Y747A-mEos2 without (16 cells) and with $Mn^{2+}$ (10 cells, 2 ind. exp.) correspond to pooled data from three independent experiments unless indicated. Where indicated, statistical significance was obtained using two-tailed, non-parametric Mann–Whitney rank sum test. Inside and outside FAs, the different conditions without $Mn^{2+}$ were compared to the corresponding β-integrin-WT condition; with $Mn^{2+}$, each given condition was compared to the value obtained without $Mn^{2+}$. Otherwise, a black line indicates which conditions were compared. The resulting P values are indicated as follows: ns: $P > 0.05$; *: $0.01 < P < 0.05$; **: $0.001 < P < 0.01$; ***: $0.0001 < P < 0.001$; ****: $P < 0.0001$.

### Figure S2

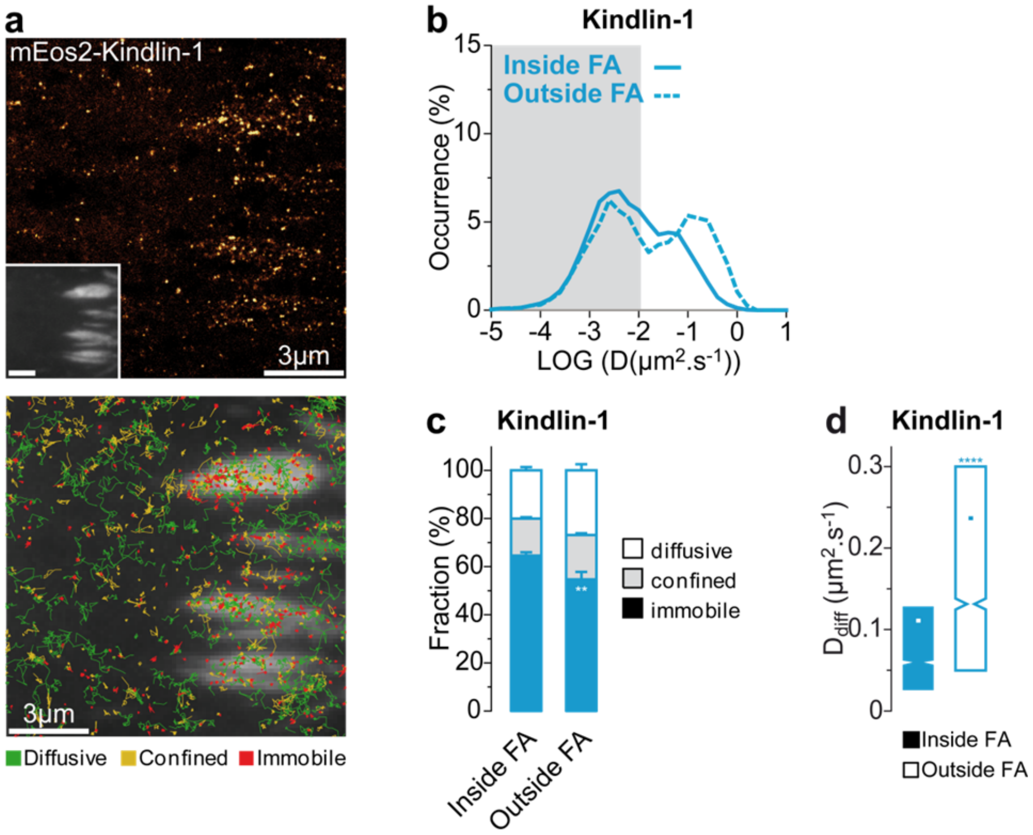


**Supplementary figure 2. Kindlin-1 undergoes lateral free diffusion along the plasma membrane.** (**a**) Top: Super-resolution PALM intensity images of mEos2-kindlin-1 in a MEF obtained from a sptPALM sequence (50 Hz, >80 s). Inset: low resolution image of GFP-paxillin, which was co-expressed for FAs labelling (scale bar: 3 µm). Bottom: color-coded trajectories overlaid on FAs labelled by GFP-paxillin (greyscale) show the diffusion modes: free diffusion (green), confined diffusion (yellow) and immobilisation (red). (**b**) Distributions of the diffusion coefficient D computed from the trajectories of mEos2-kindlin-1 obtained inside (full line) and outside (dashed line) FAs are shown in a logarithmic scale. The grey area including D values inferior to 0.011 $\mu m^2.s^{-1}$ corresponds to immobilized proteins. Values represent the average of the distributions obtained from different cells. (**c**) Fraction of mEos2-kindlin-1 undergoing free diffusion, confined diffusion or immobilisation inside (left) and outside (right) FAs. (**d**) Box plots displaying the median (notch) and mean (square) ± percentile (25–75%) of diffusion coefficients corresponding to the free diffusion trajectories of mEos2-kindlin-1 inside (left) and outside (right) FAs. Results correspond to pooled data from three independent experiments (25 cells). Where indicated, statistical significance was obtained using two-tailed, non-parametric Mann–Whitney rank sum test. The resulting P values are indicated as follows: ns: $P > 0.05$; *: $0.01 < P < 0.05$; **: $0.001 < P < 0.01$; ***: $0.0001 < P < 0.001$; ****: $P < 0.0001$.

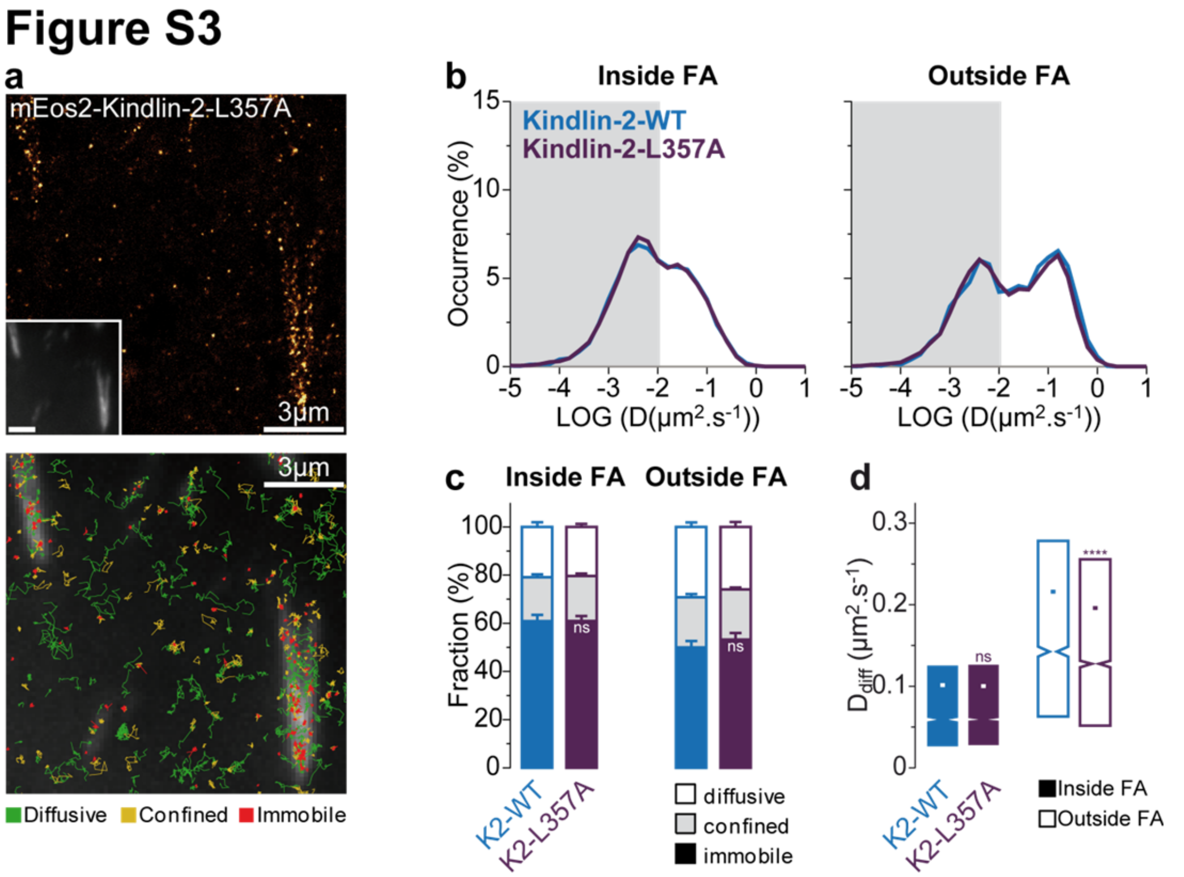


**Supplementary figure 3. Kindlin-2 membrane diffusion and immobilization does not depend on the interaction with ILK.** (**a**) Top: Super-resolution PALM intensity images of mEos2-kindlin-2-L357A in MEFs obtained from a sptPALM sequence (50 Hz, >80 s). Inset: low resolution image of GFP-paxillin, which was co-expressed for FAs labelling (scale bar: 3 µm). Bottom: color-coded trajectories overlaid on FAs labelled by GFP-paxillin (greyscale) show the diffusion modes: free diffusion (green), confined diffusion (yellow) and immobilization (red). (**b**) Distributions of the diffusion coefficient D computed from the trajectories of mEos2-kindlin-2-WT (blue) and mEos2-kindlin-2-L357A (purple) obtained inside (left) and outside FAs (right), are shown in a logarithmic scale. The grey area including D values inferior to 0.011 $\mu m^2.s^{-1}$ corresponds to immobilized proteins. Values represent the average of the distributions obtained from different cells. (**c**) Fraction of proteins undergoing free diffusion, confined diffusion or immobilization inside (left) and outside (right) adhesion sites. Values represent the average of the fractions obtained from different cells (error bars: SEM). (**d**) Box plots displaying the median (notch) and mean (square) ± percentile (25–75%) of diffusion coefficients corresponding to the free diffusion trajectories inside (left) and outside FAs (right). Results for mEos2-kindlin-2-WT (12 cells) and mEos2-kindlin-2-L357A (22 cells) correspond to pooled data from three independent experiments. Where indicated, statistical significance was obtained using two-tailed, non-parametric Mann–Whitney rank sum test. Inside and outside FAs, the mEos2-kindlin-2-L357A conditions were compared to the corresponding mEos2-kindlin-2-WT condition. The resulting P values are indicated as follows: ns: $P > 0.05$; *: $0.01 < P < 0.05$; **: $0.001 < P < 0.01$; ***: $0.0001 < P < 0.001$; ****: $P < 0.0001$.

### Figure S4

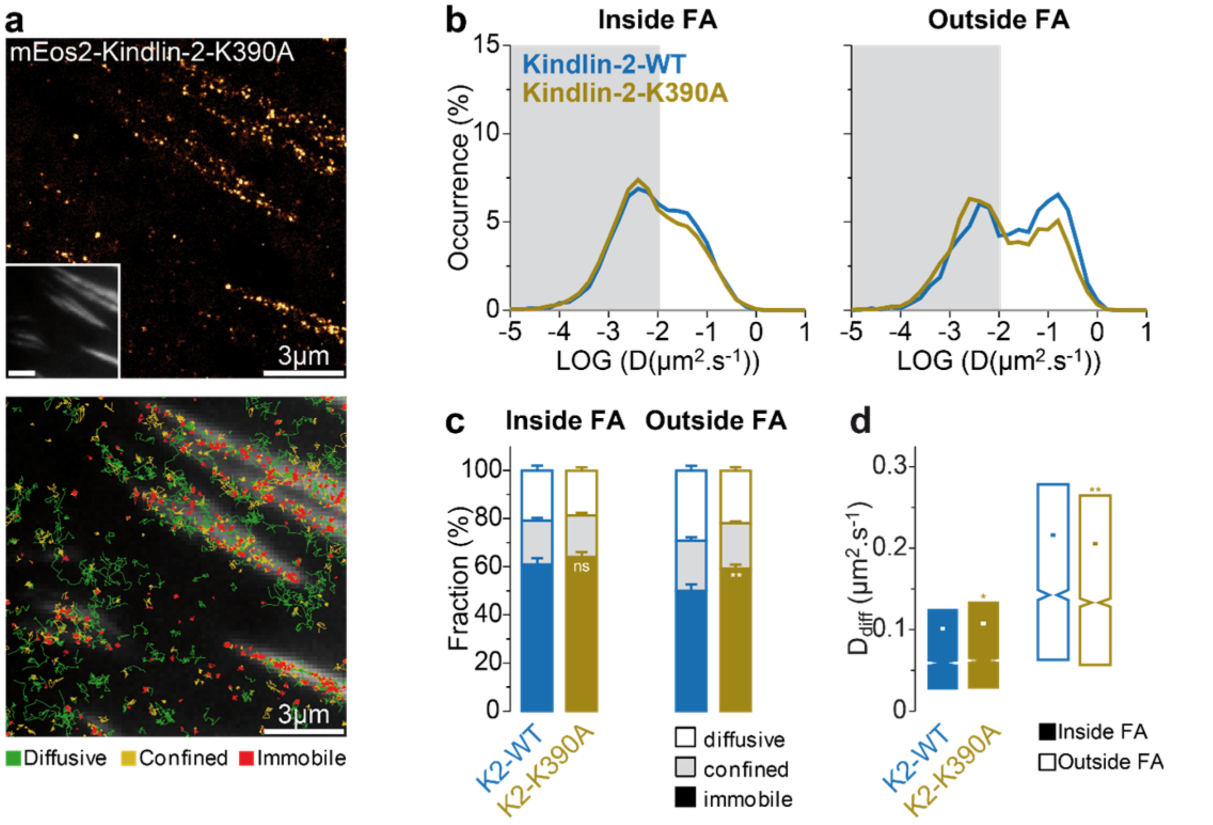


**Supplementary figure 4. Disrupting the interaction of kindlin with phosphoinositides decreases kindlin membrane diffusion.** (**a**) Top: Super-resolution PALM intensity images of mEos2-kindlin-2-K390A in MEFs obtained from a sptPALM sequence (50 Hz, >80 s). Inset: low resolution image of GFP-paxillin, which was co-expressed for FAs labelling (scale bar: 3 µm). Bottom: color-coded trajectories overlaid on FAs labelled by GFP-paxillin (greyscale) show the diffusion modes: free diffusion (green), confined diffusion (yellow) and immobilization (red). (**b**) Distributions of the diffusion coefficient D computed from the trajectories of mEos2-kindlin-2-WT (blue) and mEos2-kindlin-2-K390A (khaki) obtained inside (left) and outside FAs (right), are shown in a logarithmic scale. The grey area including D values inferior to 0.011 $\mu m^2.s^{-1}$ corresponds to immobilized proteins. Values represent the average of the distributions obtained from different cells. (**c**) Fraction of proteins undergoing free diffusion, confined diffusion or immobilization inside (left) and outside FAs (right). Values represent the average of the fractions obtained from different cells (error bars: SEM). (**d**) Box plots displaying the median (notch) and mean (square) ± percentile (25–75%) of diffusion coefficients corresponding to the free diffusion trajectories inside (left) and outside FAs (right). Results for mEos2-kindlin-2-WT (12 cells) and mEos2-kindlin-2-K390A (23 cells) correspond to pooled data from three independent experiments. Where indicated, statistical significance was obtained using two-tailed, non-parametric Mann–Whitney rank sum test. Inside and outside FAs, the mEos2-kindlin-2-K390A conditions were compared to the corresponding mEos2-kindlin-2-WT condition. The resulting P values are indicated as follows: ns: $P > 0.05$; *: $0.01 < P < 0.05$; **: $0.001 < P < 0.01$; ***: $0.0001 < P < 0.001$; ****: $P < 0.0001$.

### Figure S5

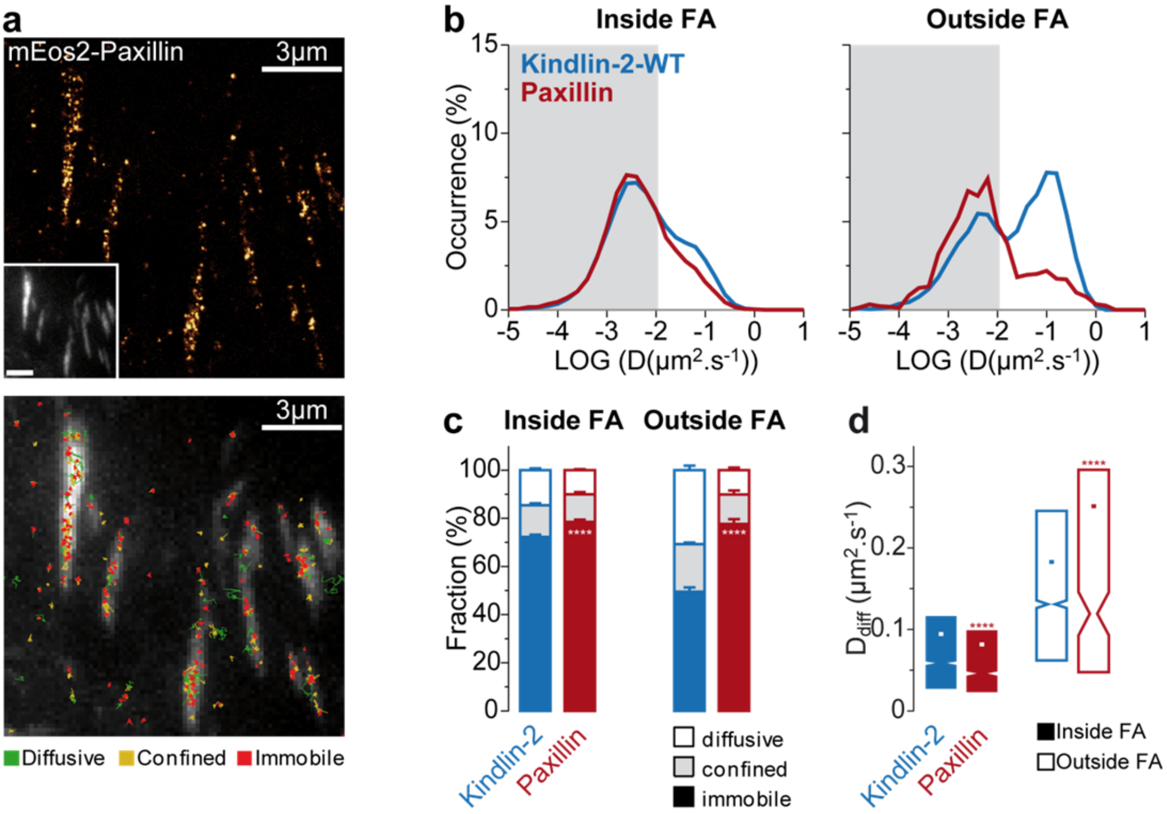


**Supplementary figure 5. Kindlin-2 diffusion is not driven by paxillin, which is mostly immobile.** (**a**) Top: Super-resolution PALM intensity images of mEos2-paxillin in MEFs obtained from a sptPALM sequence (50 Hz, >80 s). Inset: low resolution image of GFP-paxillin, which was co-expressed for FAs labelling (scale bar: 3 µm). Bottom: color-coded trajectories overlaid on FAs labelled by GFP-paxillin (greyscale) show the diffusion modes: free diffusion (green), confined diffusion (yellow) and immobilization (red). (**b**) Distributions of the diffusion coefficient D computed from the trajectories of mEos2-kindlin-2-WT (blue) and mEos2-paxillin (red) obtained inside (left) and outside FAs (right), are shown in a logarithmic scale. The grey area including D values inferior to 0.011 $\mu m^2.s^{-1}$ corresponds to immobilized proteins. Values represent the average of the distributions obtained from different cells. (**c**) Fraction of proteins undergoing free diffusion, confined diffusion or immobilization inside (left) and outside FAs (right). Values represent the average of the fractions obtained from different cells (error bars: SEM). (**d**) Box plots displaying the median (notch) and mean (square) ± percentile (25–75%) of diffusion coefficients corresponding to the free diffusion trajectories inside (left) and outside FAs (right). Results for mEos2-kindlin-2-WT (10 cells) mEos2-paxillin (12 cells) correspond to pooled data from two independent experiments. Where indicated, statistical significance was obtained using two-tailed, non-parametric Mann–Whitney rank sum test. Inside and outside FAs, the mEos2-paxillin conditions were compared to the corresponding mEos2-kindlin-2-WT condition. The resulting P values are indicated as follows: ns: $P > 0.05$; *: $0.01 < P < 0.05$; **: $0.001 < P < 0.01$; ***: $0.0001 < P < 0.001$; ****: $P < 0.0001$.

### Figure S6

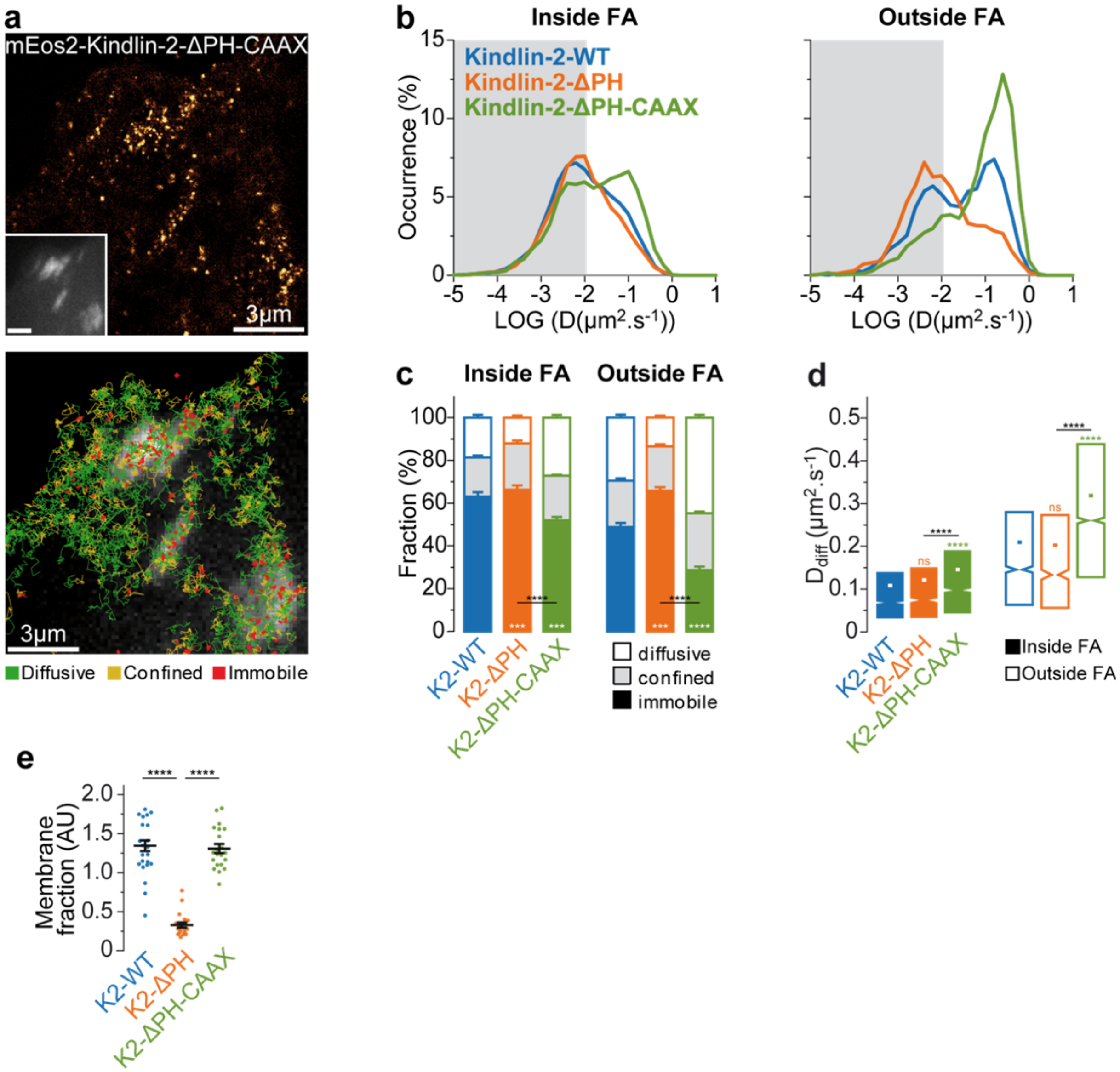


**Supplementary figure 6. Adding a CAAX sequence in kindlin-2-ΔPH induces high membrane recruitment and diffusion.** (**a**) Top: Super-resolution PALM intensity images of mEos2-kindlin-2-ΔPH-CAAX in MEFs, obtained from a sptPALM sequence (50 Hz, >80 s). Inset: low resolution image of GFP-paxillin, which was co-expressed for FAs labelling (scale bar: 3 µm). Bottom: color-coded trajectories overlaid on FAs labelled by GFP-paxillin (greyscale) show the diffusion modes: free diffusion (green), confined diffusion (yellow) and immobilization (red). (**b**) Distributions of the diffusion coefficient D computed from the trajectories of mEos2-kindlin-2-WT (blue), mEos2-kindlin-2-ΔPH (orange) and mEos2-kindlin-2-ΔPH-CAAX (green) obtained inside (left) and outside FAs (right), are shown in a logarithmic scale. The grey area including D values inferior to 0.011 $\mu m^2.s^{-1}$ corresponds to immobilized proteins. Values represent the average of the distributions obtained from different cells. (**c**) Fraction of proteins undergoing free diffusion, confined diffusion or immobilization inside (left) and outside FAs (right). Values represent the average of the fractions obtained from different cells (error bars: SEM). (**d**) Box plots displaying the median (notch) and mean (square) ± percentile (25–75%) of diffusion coefficients corresponding to the free diffusion trajectories inside (left) and outside FAs (right). (**e**) Fraction of proteins recruited at the membrane in kindlin-1, kindlin-2 knock-out cells

($Kind^{Ko}$) quantified by the ratio of the membrane-level fluorescence signal (TIRF) to the total fluorescence signal of the cell (epifluorescence). Fluorescence was measured in the red channel after photoconversion of the mEos2-genetically-coupled indicated proteins. Black bars represent mean and SEM. For sptPALM data, results for mEos2-kindlin-2-WT (15 cells), mEos2-kindlin-2-ΔPH (28 cells) and mEos2-kindlin-2-ΔPH-CAAX (18 cells) correspond to pooled data from four independent experiments. For membrane fraction data in $Kind^{Ko}$, results for mEos2-kindlin-2-WT (21 cells), mEos2-kindlin-2-ΔPH (20 cells) and mEos2-kindlin-2-ΔPH-CAAX (21 cells) come from a same experiment. Where indicated, statistical significance was obtained using two-tailed, non-parametric Mann–Whitney rank sum test. Inside and outside FAs, the different conditions were compared to the corresponding mEos2-kindlin-2-WT condition. The resulting P values are indicated as follows: ns: P > 0.05; *: 0.01 < P < 0.05; **: 0.001 < P < 0.01; ***: 0.0001 < P < 0.001; ****: P < 0.0001.

#### Figure S7

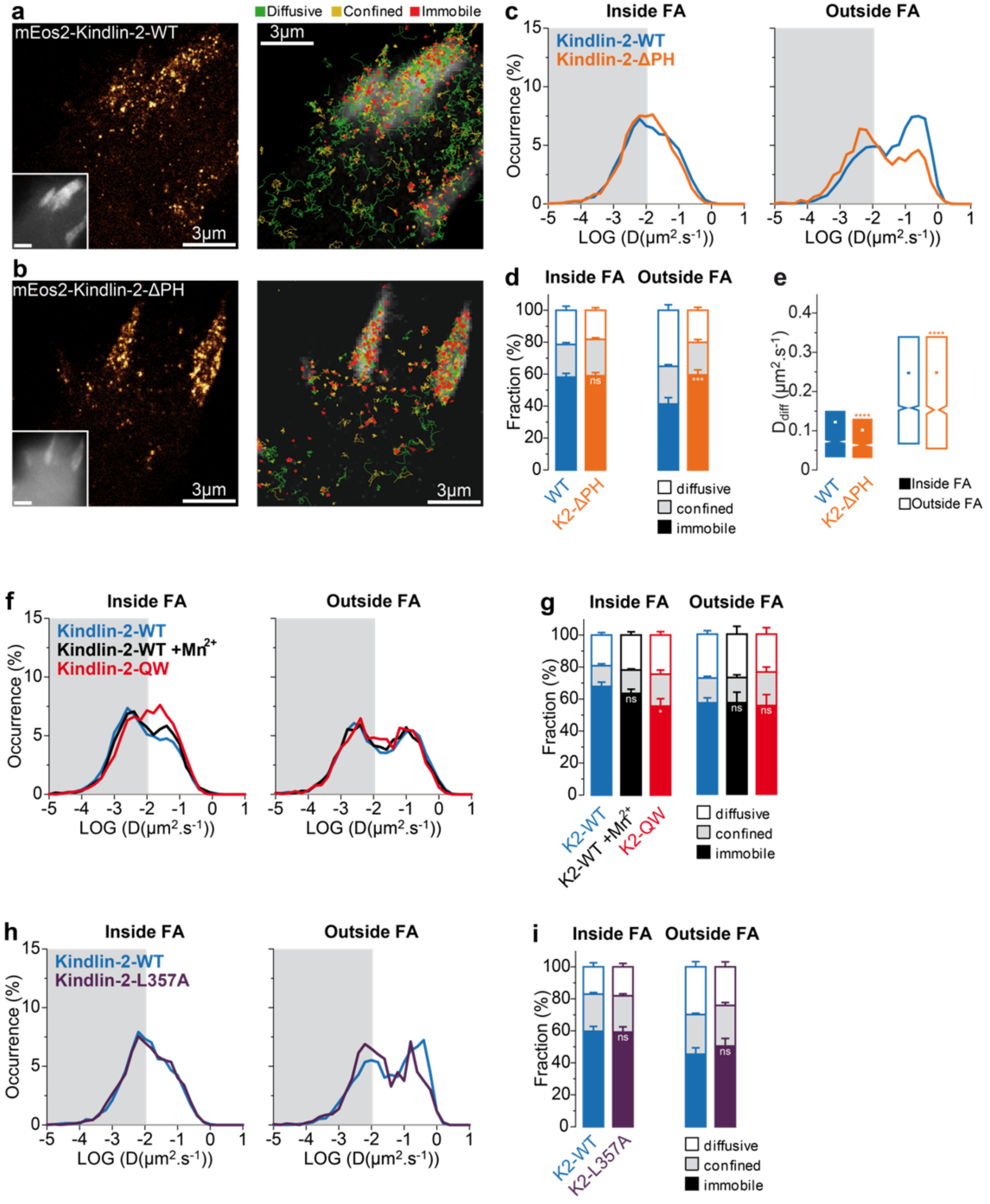


**Supplementary figure 7. MEF and Kind$^{Ko}$ (kindlin-1, kindlin-2 knock out) cells give similar sptPALM results.** All data shown were obtained in Kind$^{Ko}$ cells. (**a-b**) Left: Super-resolution intensity images of mEos2-kindlin-2-WT (a) and mEos2-kindlin-2-ΔPH (b), obtained from a sptPALM sequence (50 Hz, >80 s). Inset: low resolution image of GFP-paxillin, which was co-expressed for FAs labelling (scale bar: 3 µm). Right: color-coded trajectories overlaid on FAs labelled by GFP-paxillin (greyscale) show the diffusion modes: free diffusion (green), confined diffusion (yellow) and immobilization (red). (**c**) Distributions of the diffusion coefficient D computed from the trajectories of mEos2-kindlin-2-WT (blue) and mEos2-kindlin-2-ΔPH (orange) obtained inside (left) and outside FAs (right), are shown in a logarithmic scale. The grey area including D values inferior to 0.011 $\mu m^2.s^{-1}$ corresponds to immobilized proteins. Values represent the average of the distributions obtained from different cells. (**d**) Fraction of proteins undergoing free diffusion, confined diffusion or immobilization inside (left) and outside FAs (right). Values represent the average of the fractions obtained from different cells (error bars: SEM). (**e**) Box plots displaying the median (notch) and mean (square) ± percentile (25–75%) of diffusion coefficients corresponding to the free diffusion trajectories inside (left) and outside FAs (right). (**f**-**g**) Same as in **c**-**d**, but for mEos2-kindlin-2-WT (blue), mEos2-kindlin-2-WT in $Mn^{2+}$-stimulated cells (black) and mEos2-kindlin-2-QW614/615AA (red). (**h**-**i**) Same as in **c**-**d**, but for mEos2-kindlin-2-WT (blue) and mEos2-kindlin-2-L357A (purple). Results in (a-e) for mEos2-kindlin-2-WT (12 cells) and mEos2-kindlin-2-ΔPH (18 cells) correspond to pooled data from two independent experiments. (f,g): Results for mEos2-kindlin-2-WT (8 cells), mEos2-kindlin-2-WT +$Mn^{2+}$ (4 cells), mEos2-kindlin-2-QW614/615AA (4 cells) come from a single experiment. (h,i): Results for mEos2-kindlin-2-WT (6 cells), and mEos2-kindlin-2-L357A (4 cells) come from a single experiment. Where indicated, statistical significance was obtained using two-tailed, non-parametric Mann–Whitney rank sum test. Inside and outside FAs, the different conditions were compared to the corresponding mEos2-kindlin-2-WT condition. The resulting *P* values are indicated as follows: ns: $P > 0.05$; *: $0.01 < P < 0.05$; **: $0.001 < P < 0.01$; ***: $0.0001 < P < 0.001$; ****: $P < 0.0001$.

### Figure S8

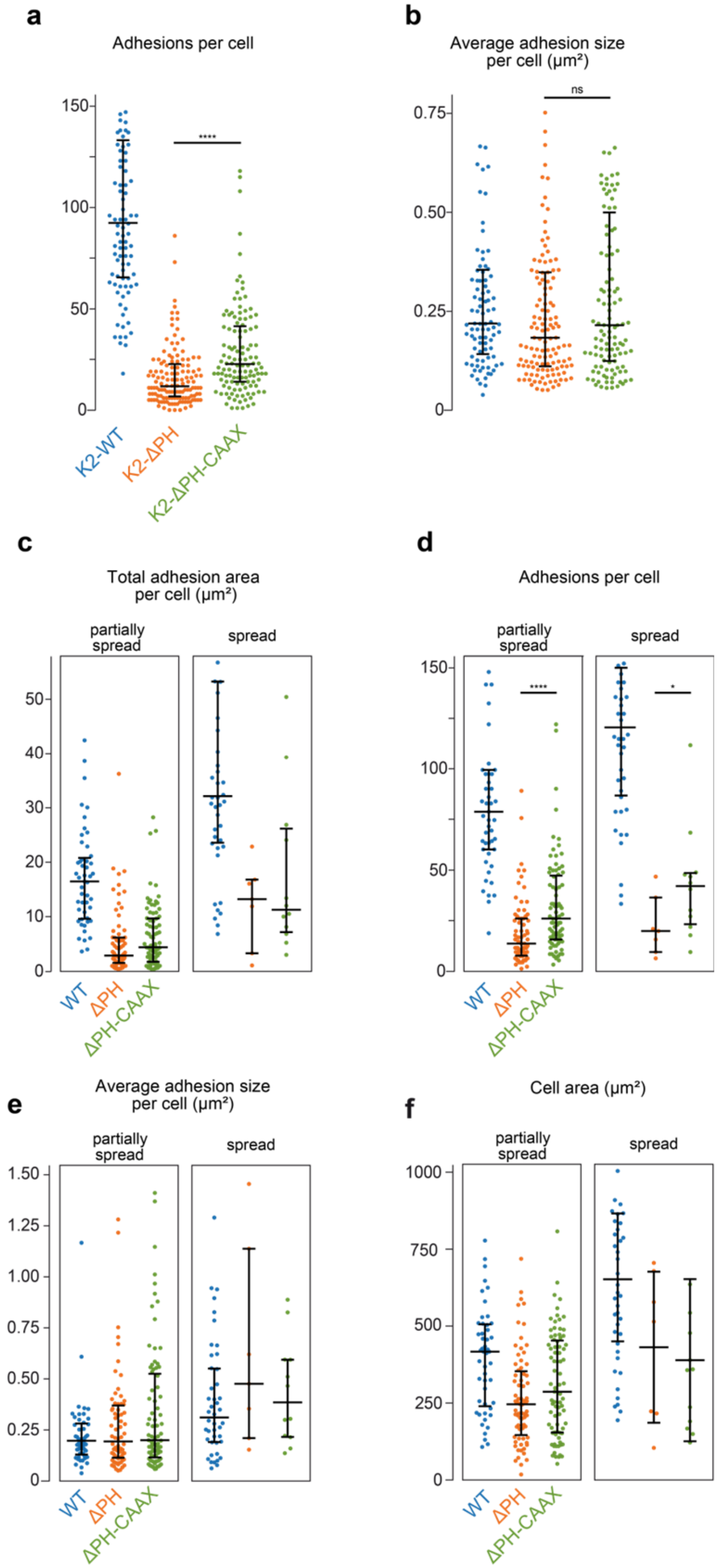

a
Adhesions per cell
****
K2-WT
K2-ΔPH
K2-ΔPH-CAAX
b
Average adhesion size per cell (μm²)
ns
c
Total adhesion area per cell (μm²)
partially spread
spread
WT
ΔPH
ΔPH-CAAX
d
Adhesions per cell
partially spread
spread
****
*
e
Average adhesion size per cell (μm²)
partially spread
spread
f
Cell area (μm²)
partially spread
spread

**Supplementary figure 8. Adhesion site measurements in function of spreading state**. Quantification of FAs number (**a**) and mean FA area (**b**) of Kind$^{Ko}$ cells 2 days after re-expression of mEos2-kindlin-2-WT (blue, 93 cells), mEos2-kindlin-2-ΔPH (orange, 134 cells) or mEos2-kindlin-2-ΔPH-CAAX (green, 120 cells) and 4h after seeding on fibronectin. Partially spread and spread cells were sorted as in Fig. 7 to generate distinct distributions of total FAs area (**c**), FAs count (**d**), mean FAs area (**e**) and cell area (**f**) for these 2 categories. Each point in the distribution represents the value obtained from a single cell. FAs were drawn manually and cell boundaries were determined by manually setting a threshold on the pixel intensity values using the TIRF GFP-paxillin images as shown in Fig. 7c. Black bars represent medians and interquartile ranges. GFP-paxillin + mEos2-kindlin-2-WT: 49 partially spread cells, 44 spread cells; GFP-paxillin + mEos2-kindlin-2-ΔPH: 72 partially spread cells, 7 spread cells; GFP-paxillin + mEos2-kindlin-2-ΔPH-CAAX: 81 partially spread cells, 12 spread cells; the results correspond to pooled data from four independent experiments. Where indicated, statistical significance was obtained using two-tailed, non-parametric Mann–Whitney rank sum test. The resulting P values are indicated as follows: ns: $P > 0.05$; *: $0.01 < P < 0.05$; **: $0.001 < P < 0.01$; ***: $0.0001 < P < 0.001$; ****: $P < 0.0001$.

#### SUPPLEMENTARY TABLES

**Table S1.** Results of sptPALM experiments (FAs: Focal adhesions).

| | β1-integrin-WT | β1-integrin-Y795A | β1-integrin-Y783A | β1-integrin-WT + Mn2+ | β1-integrin-Y795A +Mn2+ |
|---|---|---|---|---|---|
| cells | 16 | 22 | 16 | 15 | 22 |
| trajectories inside FAs | 25270 | 12531 | 11606 | 20775 | 11243 |
| trajectories outside FAs | 64523 | 55527 | 74583 | 56565 | 66450 |
| | **Mean ± SEM (median)** | **Mean ± SEM (median)** | **Mean ± SEM (median)** | **Mean ± SEM (median)** | **Mean ± SEM (median)** |
| immobile inside FAs (%) | 71.7 ± 1.5 (72.6) | 47.9 ± 2.9 (47.7) | 41.2 ± 1.7 (41.2) | 83.4 ± 1.1 (83.9) | 60.3 ± 2.1 (60.3) |
| confined inside FAs (%) | 11.2 ± 0.5 (11.3) | 20.5 ± 1.1 (20.8) | 22.6 ± 0.8 (21.8) | 7.1 ± 0.7 (7.1) | 14.8 ± 0.8 (15.3) |
| diffusive inside FAs (%) | 17.1 ± 1.2 (17.4) | 31.6 ± 2 (30.9) | 36.3 ± 1 (37.1) | 9.5 ± 0.6 (9.6) | 24.9 ± 1.6 (24.1) |
| immobile outside FAs (%) | 31.6 ± 2.3 (31.4) | 19.4 ± 1.1 (19.8) | 19.1 ± 1.2 (18.3) | 58.7 ± 2.6 (57.3) | 36.8 ± 2.5 (35.9) |
| confined outside FAs (%) | 20 ± 0.6 (20.3) | 25.5 ± 0.5 (25.7) | 22.8 ± 0.4 (23.1) | 13.4 ± 1 (13.6) | 21.1 ± 1.1 (20.8) |
| diffusive outside FAs (%) | 48.3 ± 2 (48) | 55.1 ± 1.1 (55) | 58.1 ± 1.2 (58.1) | 27.9 ± 2 (29.1) | 42 ± 1.8 (41.9) |
| $D_{diff}$ inside x$10^3$ ($\mu m^2.s^{-1}$) | 195.9 ± 3.4 (127.5) | 237.5 ± 4 (159.3) | 217.7 ± 3 (162.1) | 154.8 ± 3.9 (93.7) | 224.4 ± 4.2 (162.8) |
| $D_{diff}$ outside x$10^3$ ($\mu m^2.s^{-1}$) | 330.7 ± 1.3 (293.6) | 332.5 ± 1.3 (293.1) | 342 ± 1 (312.4) | 275.4 ± 1.7 (236.5) | 318.7 ± 1.4 (275.5) |

| | β1-integrin-Y783A +Mn2+ | β3-integrin-WT | β3-integrin-Y759A | β3-integrin-Y747A | β3-integrin-WT +Mn2+ |
|---|---|---|---|---|---|
| cells | 20 | 16 | 20 | 16 | 13 |
| trajectories inside FA | 13615 | 11347 | 18438 | 10624 | 16487 |
| trajectories outside FA | 82547 | 9355 | 34797 | 28579 | 78006 |
| | **Mean ± SEM (median)** | **Mean ± SEM (median)** | **Mean ± SEM (median)** | **Mean ± SEM (median)** | **Mean ± SEM (median)** |
| immobile inside FA (%) | 68 ± 2.1 (69.7) | 61.7 ± 1.5 (62.2) | 57 ± 1.5 (58.1) | 46.1 ± 2.5 (45.7) | 82.7 ± 1.5 (85.9) |
| confined inside FA (%) | 12.1 ± 0.8 (12.3) | 20.2 ± 1.1 (20.1) | 18.4 ± 0.8 (17.3) | 19.5 ± 0.7 (19.5) | 13.1 ± 1.1 (14.7) |
| diffusive inside FA (%) | 19.9 ± 1.4 (18.5) | 18.1 ± 1.4 (17.6) | 24.6 ± 1.3 (24.3) | 34.4 ± 2.2 (34.4) | 6.3 ± 0.8 (4.9) |
| immobile outside FA (%) | 51.5 ± 2.3 (51.4) | 49 ± 2.9 (49) | 41 ± 2.5 (40.6) | 36.3 ± 2.3 (37.2) | 77 ± 1.7 (78.9) |
| confined outside FA (%) | 15.1 ± 0.6 (15) | 19.3 ± 1.2 (19.8) | 20.8 ± 0.9 (21.7) | 20.1 ± 0.8 (19.9) | 15.2 ± 1.2 (13.9) |
| diffusive outside FA (%) | 33.4 ± 1.8 (32.6) | 31.6 ± 2.4 (30.3) | 38.2 ± 1.9 (37.9) | 43.7 ± 2 (42.6) | 11.7 ± 1.4 (10) |

| | | | | | |
|---|---|---|---|---|---|
| $D_{diff}$ inside x$10^3$ (µm².$s^{-1}$) | 205.2 ± 4 (144.3) | 119.6 ± 2.7 (78.5) | 150.2 ± 2.1 (109.1) | 162.5 ± 2.4 (126.5) | 95.6 ± 3.2 (53.6) |
| $D_{diff}$ outside x$10^3$ (µm².$s^{-1}$) | 284.9 ± 1.3 (245.8) | 257 ± 3.6 (213.1) | 242.5 ± 1.6 (202.5) | 246.6 ± 1.7 (208.6) | 199.4 ± 3.7 (156.7) |

| | β3-integrin-Y759A +Mn2+ | β3-integrin-Y747A +Mn2+ | Kindlin-2-WT (Fig 2) | Talin-1 | Kindlin-1 |
|---|---|---|---|---|---|
| cells | 11 | 10 | 13 | 8 | 25 |
| trajectories inside FA | 15660 | 12783 | 15475 | 18159 | 17305 |
| trajectories outside FA | 18685 | 24649 | 9792 | 7940 | 17940 |
| | Mean ± SEM (median) | Mean ± SEM (median) | Mean ± SEM (median) | Mean ± SEM (median) | Mean ± SEM (median) |
| immobile inside FA (%) | 80.4 ± 0.5 (80) | 80.2 ± 1.4 (80.3) | 64.9 ± 2.3 (67.4) | 81.6 ± 1.5 (80.8) | 64.6 ± 1.4 (62.9) |
| confined inside FA (%) | 12.1 ± 0.7 (12) | 11 ± 0.9 (9.2) | 17.1 ± 0.8 (16.3) | 11.9 ± 0.8 (11.7) | 15.4 ± 0.6 (14.4) |
| diffusive inside FA (%) | 6.5 ± 0.8 (5.2) | 7.7 ± 1.5 (6.9) | 18 ± 1.6 (16.9) | 6.5 ± 0.9 (5.7) | 20 ± 1.4 (20.5) |
| immobile outside FA (%) | 76.1 ± 2 (76.8) | 74.5 ± 1.6 (76.1) | 52.8 ± 2.8 (52.8) | 73.2 ± 3.4 (74) | 54.7 ± 3.1 (53.5) |
| confined outside FA (%) | 15.2 ± 1 (15.6) | 11.3 ± 0.5 (11.1) | 20.2 ± 0.9 (20.5) | 14.3 ± 1.5 (15.4) | 18.5 ± 0.8 (17.6) |
| diffusive outside FA (%) | 8.8 ± 1.2 (7.3) | 10.3 ± 1.9 (8.9) | 27 ± 2.1 (28) | 12.6 ± 2.2 (10.2) | 26.9 ± 2.5 (30.6) |
| $D_{diff}$ inside x$10^3$ (µm².$s^{-1}$) | 105.5 ± 5.4 (64.4) | 107.7 ± 3.4 (72.1) | 96.5 ± 2.1 (58.2) | 114.7 ± 4.6 (54.7) | 110.7 ± 2.7 (59.6) |
| $D_{diff}$ outside x$10^3$ (µm².$s^{-1}$) | 200.5 ± 7.9 (149.1) | 169.9 ± 3.8 (121.3) | 175.7 ± 3.9 (111.8) | 120.6 ± 4.8 (75.3) | 236.6 ± 5 (131.7) |

| | Kindlin-2-WT (Fig 3) | Kindlin-2-WT + Mn2+ activation | Kindlin-2-QW | Kindlin-2-WT (Fig S3,4) | Kindlin-2-L357A |
|---|---|---|---|---|---|
| cells | 17 | 17 | 33 | 12 | 22 |
| trajectories inside FA | 24941 | 33799 | 38495 | 15959 | 39848 |
| trajectories outside FA | 10677 | 42414 | 17831 | 7926 | 16832 |
| | Mean ± SEM (median) | Mean ± SEM (median) | Mean ± SEM (median) | Mean ± SEM (median) | Mean ± SEM (median) |
| immobile inside FA (%) | 67.1 ± 2.3 (67.6) | 65.8 ± 2 (63) | 53.3 ± 1.8 (55.4) | 60.8 ± 2.7 (61.2) | 60.9 ± 2.1 (60.5) |
| confined inside FA (%) | 14.6 ± 1.2 (12.8) | 14.6 ± 0.7 (15) | 19.8 ± 0.9 (18.7) | 18.3 ± 1.2 (18.1) | 18.8 ± 1 (18.1) |
| diffusive inside FA (%) | 18.3 ± 1.3 (18.9) | 19.6 ± 1.5 (20.7) | 26.9 ± 1.2 (26.2) | 20.9 ± 1.9 (19.5) | 20.3 ± 1.3 (20.1) |
| immobile outside FA (%) | 58.9 ± 2.5 (59.7) | 58.1 ± 2.5 (57.2) | 52.8 ± 2.6 (50.2) | 49.9 ± 2.6 (49.5) | 53.2 ± 2.8 (53) |
| confined outside FA (%) | 17.1 ± 0.8 (17.3) | 15.8 ± 0.9 (15.3) | 19.7 ± 1 (21) | 20.9 ± 1.3 (20.7) | 20.8 ± 0.8 (20.7) |
| diffusive outside FA (%) | 24 ± 2 (21.7) | 26.1 ± 1.8 (26.2) | 27.6 ± 1.9 (28.9) | 29.2 ± 1.9 (28.9) | 26 ± 2.1 (26.7) |

| | | | | | |
|---|---|---|---|---|---|
| $D_{diff}$ inside x$10^3$ ($\mu m^2.s^{-1}$) | 76.1 ± 1.6 (41.2) | 92.1 ± 1.5 (50.9) | 102 ± 1.3 (58.2) | 101.7 ± 2 (58.9) | 100 ± 1.2 (59.3) |
| $D_{diff}$ outside x$10^3$ ($\mu m^2.s^{-1}$) | 43.8 ± 1.4 (0) | 190 ± 2.2 (114.7) | 190.6 ± 3.4 (98.1) | 215.9 ± 4.2 (142.6) | 196 ± 2.9 (127.3) |

| | Kindlin-2-K390A | Kindlin-2-WT (fig 4, S6) | Kindlin-2-ΔPH (MEF) | Kindlin-2-ΔPH-CAAX | PH domain |
|---|---|---|---|---|---|
| cells | 23 | 15 | 28 | 18 | 18 |
| trajectories inside FA | 39632 | 16955 | 11295 | 10926 | 5831 |
| trajectories outside FA | 18734 | 6891 | 6483 | 11829 | 10397 |
| | **Mean ± SEM (median)** | **Mean ± SEM (median)** | **Mean ± SEM (median)** | **Mean ± SEM (median)** | **Mean ± SEM (median)** |
| immobile inside FA (%) | 64.1 ± 2.1 (68) | 63 ± 2.1 (64.3) | 66.3 ± 2.1 (66.3) | 52 ± 1.5 (50.5) | 60.6 ± 2.8 (57.3) |
| confined inside FA (%) | 17.3 ± 1 (14.9) | 18.4 ± 0.9 (18.3) | 21.6 ± 1.3 (21) | 20.8 ± 0.5 (21.4) | 17.4 ± 1 (17.8) |
| diffusive inside FA (%) | 18.7 ± 1.2 (16.7) | 18.6 ± 1.3 (18.5) | 12.1 ± 1 (12.3) | 27.2 ± 1.3 (27.3) | 22 ± 2.1 (24.7) |
| immobile outside FA (%) | 59.2 ± 1.7 (59.2) | 48.8 ± 1.9 (49.6) | 65.7 ± 1.7 (68.6) | 28.7 ± 1.7 (28.5) | 49.1 ± 4.3 (42.3) |
| confined outside FA (%) | 19 ± 0.6 (18.7) | 21.7 ± 1.1 (21) | 20.8 ± 1 (19.8) | 26.6 ± 0.8 (26.9) | 19.2 ± 1.4 (18.9) |
| diffusive outside FA (%) | 21.8 ± 1.4 (22.9) | 29.5 ± 1.4 (29.9) | 13.5 ± 0.9 (13) | 44.7 ± 1.3 (45.1) | 31.7 ± 3.1 (34.9) |
| $D_{diff}$ inside x$10^3$ ($\mu m^2.s^{-1}$) | 107.5 ± 1.5 (62.2) | 108.5 ± 2.2 (68.1) | 121.8 ± 3.9 (74.3) | 146.2 ± 2.8 (97.6) | 241.3 ± 9.2 (141.5) |
| $D_{diff}$ outside x$10^3$ ($\mu m^2.s^{-1}$) | 205.2 ± 3.5 (133.2) | 209.6 ± 4.7 (146) | 202.3 ± 7.4 (134) | 319 ± 3.5 (260.4) | 343.9 ± 6.6 (242) |

| | Kindlin-2-WT (fig S5) | Paxillin | Kindlin-2-WT (Kind$^{Ko}$, fig S7a-e) | Kindlin-2-ΔPH (Kind$^{Ko}$) | Kindlin-2-WT (Kind$^{Ko}$, fig S7f,g) |
|---|---|---|---|---|---|
| cells | 10 | 12 | 12 | 18 | 8 |
| trajectories inside FA | 22501 | 16678 | 14940 | 9318 | 14279 |
| trajectories outside FA | 11406 | 2193 | 15985 | 8634 | 6115 |
| | **Mean ± SEM (median)** | **Mean ± SEM (median)** | **Mean ± SEM (median)** | **Mean ± SEM (median)** | **Mean ± SEM (median)** |
| immobile inside FA (%) | 72.2 ± 1 (73) | 78.5 ± 0.9 (78.9) | 56.9 ± 2.5 (58.7) | 58.9 ± 2 (59.7) | 67.9 ± 2.7 (68.9) |
| confined inside FA (%) | 13.2 ± 0.8 (12.9) | 11.4 ± 0.9 (11.4) | 20.5 ± 1.1 (19.1) | 22.8 ± 1 (23) | 13 ± 1.2 (12.1) |
| diffusive inside FA (%) | 14.6 ± 0.8 (15.1) | 10.1 ± 0.4 (10.2) | 22.3 ± 2.5 (20.7) | 18.2 ± 1.6 (18.5) | 19.1 ± 1.6 (19.3) |
| immobile outside FA (%) | 49.5 ± 1.8 (49.8) | 77.6 ± 2 (79.9) | 59.5 ± 3.2 (61.9) | 20.1 ± 1.8 (17.8) | 57.1 ± 3.1 (59.5) |
| confined outside FA (%) | 19.8 ± 0.7 (20.3) | 12.3 ± 1.7 (9.4) | 23.6 ± 1 (24.3) | 20.4 ± 1.8 (19.9) | 15.6 ± 1 (14.7) |
| diffusive outside FA (%) | 30.8 ± 1.8 (29.7) | 10.1 ± 1 (10.4) | 40.5 ± 3.7 (40.6) | 35.8 ± 3.2 (35.3) | 27.4 ± 2.2 (26.3) |
| $D_{diff}$ inside x$10^3$ ($\mu m^2.s^{-1}$) | 106 ± 3.1 (59.9) | 87 ± 1.9 (53.8) | 126.8 ± 2.4 (74) | 102.7 ± 2.6 (63.8) | 103 ± 2.8 (58.9) |

| | | | | | |
|---|---|---|---|---|---|
| $D_{diff}$ outside x10$^3$ (µm².s$^{-1}$) | 202.9 ± 6.4 (117.3) | 226.8 ± 19.8 (92.6) | 275.7 ± 3.6 (176.3) | 249.5 ± 6.7 (153.9) | 270.1 ± 8.1 (160.3) |

| | Kindlin-2-WT + Mn2+ activation (Kind$^{Ko}$) | Kindlin-2-QW (Kind$^{Ko}$) | Kindlin-2-WT (Kind$^{Ko}$, fig S7h,i) | Kindlin-2-L357A (Kind$^{Ko}$) |
|---|---|---|---|---|
| cells | 4 | 4 | 6 | 4 |
| trajectories inside FA | 11114 | 2798 | 6095 | 3033 |
| trajectories outside FA | 9745 | 2535 | 2895 | 1297 |
| | **Mean ± SEM (median)** | **Mean ± SEM (median)** | **Mean ± SEM (median)** | **Mean ± SEM (median)** |
| immobile inside FA (%) | 63.6 ± 2.6 (63.4) | 55.6 ± 4.6 (54.8) | 59.7 ± 3.1 (63) | 59.2 ± 3.2 (62) |
| confined inside FA (%) | 14.6 ± 0.7 (14.7) | 20 ± 2.6 (20.9) | 23.2 ± 1 (23.9) | 22.7 ± 1.3 (21.8) |
| diffusive inside FA (%) | 21.8 ± 2.1 (22.2) | 24.4 ± 2.2 (24.3) | 17.1 ± 2.5 (15.6) | 18.1 ± 2.1 (16.7) |
| immobile outside FA (%) | 57.2 ± 6.6 (56.8) | 55.4 ± 6.9 (53.6) | 45.4 ± 4 (43.5) | 50.5 ± 4.7 (50.9) |
| confined outside FA (%) | 15.7 ± 1.7 (16.1) | 21 ± 3.1 (20.6) | 24.8 ± 0.8 (25.5) | 25.4 ± 1.8 (24.8) |
| diffusive outside FA (%) | 27.1 ± 4.9 (26.9) | 23.6 ± 4.1 (24.8) | 29.8 ± 3.2 (30.6) | 24.1 ± 3.1 (23) |
| $D_{diff}$ inside x10$^3$ (µm².s$^{-1}$) | 103.4 ± 2.4 (62.1) | 90.2 ± 3.3 (58.3) | 132.9 ± 4.7 (82.3) | 135.7 ± 5.6 (92.6) |
| $D_{diff}$ outside x10$^3$ (µm².s$^{-1}$) | 264.1 ± 5.9 (148.5) | 242 ± 10.8 (138.7) | 333.2 ± 10.3 (237.1) | 289.9 ± 18 (195.4) |

**Table S2.** Results of immobilization time measurements inside focal adhesions by PALM.

| | β1-integrin-WT | β1-integrin-Y795A | β1-integrin-Y783A | β3-integrin-WT | β3-integrin-Y759A | β3-integrin-Y747A |
|---|---|---|---|---|---|---|
| cells | 3 | 6 | 6 | 6 | 5 | 5 |
| immobilizations | 296 | 229 | 272 | 557 | 496 | 464 |
| Immobilization time (mean ± SEM (median), seconds) | 22.29 ± 1.62 (12) | 10.81 ± 1.12 (5) | 15.28 ± 1.72 (5.25) | 40.99 ± 1.93 (25) | 32.97 ± 1.85 (20) | 31.05 ± 1.79 (19) |

**Table S3.** Results of membrane fraction experiments.

| | Kindlin-2-WT (MEF) | Kindlin-2-ΔPH (MEF) | Kindlin-2-QW | Kindlin-2-K390A | Kindlin-2-L357A |
|---|---|---|---|---|---|
| cells | 28 | 27 | 15 | 15 | 15 |
| Membrane fraction (Mean ± SEM (median), arbitrary unit) | 1.41 ± 0.05 (1.35) | 0.64 ± 0.02 (0.62) | 1.05 ± 0.03 (1.03) | 1.24 ± 0.05 (1.25) | 1.23 ± 0.05 (1.24) |

| | β3-integrin-WT | mEos2 | Kindlin-2-WT (Kind$^{Ko}$) | Kindlin-2-ΔPH (Kind$^{Ko}$) | Kindlin-2-ΔPH-CAAX |
|---|---|---|---|---|---|
| cells | 17 | 13 | 21 | 20 | 21 |
| Membrane fraction (AU, mean ± SEM (median)) | 1.32 ± 0.04 (1.31) | 0.56 ± 0.02 (0.53) | 1.35 ± 0.07 (1.33) | 0.33 ± 0.03 (0.28) | 1.31 ± 0.06 (1.25) |

**Table S4.** Results of DONALD experiments. Molecules detected above 200 nm were considered as non-specific and were excluded.

| | Paxillin | Kindlin-2-WT | Kindlin-2-QW | Kindlin-2-ΔPH | Kindlin-2-ΔPH-CAAX |
|---|---|---|---|---|---|
| cells | 6 | 6 | 9 | 7 | 7 |
| detections | 2720922 | 2112741 | 2407581 | 2572448 | 3257051 |
| $z_{median}$ (mean ± SEM (median), nm) | 73.1 ± 0.03 (67.6) | 68.1 ± 0.04 (63.0) | 71.8 ± 0.03 (66.9) | 82.7 ± 0.03 (78.7) | 60.3 ± 0.03 (53.3) |

**Table S5.** Results of focal adhesion (FA) enrichment experiments.

| | Kindlin-2-WT | mEos2 | Kindlin-2-ΔPH | Kindlin-2-QW | Kindlin-2-ΔPH-CAAX |
|---|---|---|---|---|---|
| cells | 20 | 15 | 20 | 20 | 20 |
| FA enrichment (Mean ± SEM (median), arbitrary unit) | 1.8 ± 0.07 (1.72) | 0.99 ± 0.03 (0.94) | 1.15 ± 0.03 (1.16) | 1.26 ± 0.02 (1.24) | 1.5 ± 0.03 (1.49) |

| | Kindlin-2-L357A | Kindlin-2-K390A |
|---|---|---|
| cells | 12 | 12 |
| FA enrichment (Mean ± SEM (median), arbitrary unit) | 1.46 ± 0.06 (1.47) | 1.75 ± 0.06 (1.76) |

**Table S6.** Results of cell spreading experiments. Percentages are expressed in the format "mean ± SEM (median)".

| | Paxillin | Kindlin-2-WT | Kindlin-2-QW | Kindlin-2-ΔPH | Kindlin-2-ΔPH-CAAX | Kindlin-2-WT-CAAX |
|---|---|---|---|---|---|---|
| cells | 141 | 312 | 284 | 311 | 309 | 289 |
| non-spread (%) | 96.2 ± 3 (98.3) | 21.8 ± 4.6 (23) | 35.4 ± 6.9 (41.8) | 72.6 ± 7.3 (78.8) | 43.5 ± 3.1 (44.6) | 27.3 ± 7.9 (21.8) |
| partially spread (%) | 3.8 ± 3 (1.7) | 57.7 ± 1.7 (58.9) | 56 ± 4.4 (52.4) | 24.6 ± 6.4 (19.2) | 47.8 ± 4.3 (45.5) | 54.1 ± 1.9 (53.5) |
| spread (%) | 0 ± 0 (0) | 20.5 ± 6.1 (17) | 8.5 ± 2.7 (7.1) | 2.9 ± 0.9 (2) | 8.7 ± 1.3 (9.9) | 18.6 ± 6.3 (24.8) |

**Table S7.** Results of focal adhesion experiments. Except cell numbers, values are expressed in the format "mean ± SEM (median)". FA: Focal adhesion.

| | Kindlin-2-WT | Kindlin-2- ΔPH | Kindlin-2- ΔPH-CAAX |
|---|---|---|---|
| cells | 93 | 134 | 120 |
| cell area (μm²) | 542.6 ± 37.1 (475) | 205.9 ± 13.9 (165.6) | 272.2 ± 17.1 (228.4) |
| FA total area (μm²) | 29.6 ± 2.8 (22.5) | 4.3 ± 0.5 (2.4) | 7.6 ± 0.7 (5.4) |
| FA count | 106.2 ± 6.6 (92) | 16 ± 1.2 (11) | 28.3 ± 2 (22) |
| Average adhesion size / cell (μm²) | 0.305 ± 0.025 (0.219) | 0.276 ± 0.024 (0.183) | 0.326 ± 0.026 (0.215) |

| | Kindlin-2-WT (partially spread) | Kindlin-2- ΔPH (partially spread) | Kindlin-2- ΔPH-CAAX (partially spread) |
|---|---|---|---|
| cells | 49 | 72 | 81 |
| cell area (μm²) | 391.1 ± 23.9 (416.9) | 265.4 ± 17.6 (246.2) | 310.5 ± 18.8 (286.6) |
| FA total area (μm²) | 16.9 ± 1.3 (16.3) | 5 ± 0.7 (2.7) | 6.2 ± 0.7 (4.2) |
| FA count | 88.6 ± 8.2 (76) | 18.5 ± 1.9 (13) | 31 ± 2.5 (25) |
| Average adhesion size / cell (μm²) | 0.227 ± 0.024 (0.197) | 0.288 ± 0.033 (0.194) | 0.336 ± 0.034 (0.2) |

| | Kindlin-2-WT (spread) | Kindlin-2- ΔPH (spread) | Kindlin-2- ΔPH-CAAX (spread) |
|---|---|---|---|
| cells | 44 | 7 | 12 |
| cell area (μm²) | 710.1 ± 64.5 (652.2) | 431.5 ± 92.7 (514.4) | 389.4 ± 76.1 (358.5) |
| FA total area (μm²) | 43.8 ± 5 (32) | 12 ± 2.9 (13.1) | 17.4 ± 4.3 (11.1) |
| FA count | 125.8 ± 9.6 (116.5) | 21.3 ± 5.3 (19) | 41.3 ± 7.6 (40.5) |
| Average adhesion size / cell (μm²) | 0.391 ± 0.043 (0.311) | 0.629 ± 0.185 (0.476) | 0.434 ± 0.073 (0.385) |

#### SUPPLEMENTARY MOVIE LEGENDS

**Supplementary Movie 1. Kindlin-2 undergoes free diffusion along the plasma membrane inside and outside FAs contrary to talin.**

This movie shows the time course of single protein tracking experiments depicted in Fig. 2. It covers a period of about 20 seconds, during which single kindlin-2-WT (left) and talin (right) proteins labeled by mEos2 are tracked inside and outside FAs (revealed by GFP-paxillin and outlined by blue lines). Scale bar = 1.6 µm. Speed, 0.6X.

**Supplementary Movie 2. Deletion of the PH domain (kindlin-2-ΔPH) strongly inhibits membrane free diffusion both inside and outside FAs.**

This movie shows the time course of the single protein tracking experiment depicted in Fig. 4. It covers a period of about 20 seconds, during which single kindlin-2-ΔPH proteins labeled by mEos2 are tracked inside and outside FAs (revealed by GFP-paxillin and outlined by blue lines). Scale bar = 1.6 µm. Speed, 0.6X.

**Supplementary Movie 3. Restoring membrane association of kindlin-2-ΔPH by adding a CAAX prenylation sequence induces membrane recruitment and diffusion inside and outside FAs.**

This movie shows the time course of the single protein tracking experiment depicted in Supplementary Fig. S6. It covers a period of about 20 seconds, during which single kindlin-2-ΔPH-CAAX proteins labeled by mEos2 are tracked inside and outside FAs (revealed by GFP-paxillin and outlined by blue lines). Scale bar = 1.6 µm. Speed, 0.6X.
